# Spatiotemporal trends in polychlorinated biphenyl concentrations in seafood based on long-term monitoring and remediation in New Bedford Harbor, Massachusetts

**DOI:** 10.1101/675355

**Authors:** Kathryn A. Crawford, Jennifer J. Schlezinger, Paul Craffey, Wendy Heiger-Bernays

**Affiliations:** Department of Environmental Health, Boston University School of Public Health, Boston, MA, USA; Response and Remediation Division, Bureau of Waste Site Cleanup, Massachusetts Department of Environmental Protection, Boston, MA, USA

**Keywords:** Polychlorinated biphenyls, PCBs, bioaccumulation, seafood consumption, risk assessment, scup, clam

## Abstract

Electrical manufacturing near New Bedford Harbor (NBH), MA in the mid-1900s led to severe polychlorinated biphenyl (PCB) contamination, which resulted in the harbor’s designation as a Superfund Site. Restrictions on the harvest of seafood from NBH have been in effect since 1979. Efforts to reduce the overall mass of PCBs in NBH by dredging PCB-contaminated sediments have been ongoing since the late 1980s. One goal of dredging is to reduce PCB concentrations in NBH seafood, monitored annually by Massachusetts Department of Environmental Protection (DEP) and Massachusetts Division of Marine Fisheries (MDMF). We used PCB concentrations in quahogs (2003-2016) and scup (2003-2014) to evaluate PCB distribution across seafood management areas and over time. Seafood PCB concentrations were used to evaluate improvements in environmental quality by examining total PCBs and patterns of PCB congeners within homolog groups, and by assessing human cancer risk from seafood consumption in the past (1980) and present (2012-2016). PCB concentrations in quahogs generally declined with increased time and distance from the PCB source, as does the cancer risk associated with their consumption. PCB concentrations in scup follow similar spatial patterns but show high annual variability. We conclude that quahogs are a reliable proxy for *in-situ* conditions, environmental quality, and human health risk.

## 1. Introduction

The New Bedford Harbor (NBH) region of Massachusetts was a commercial epicenter for the northeastern United States during the industrial and manufacturing revolutions. Today, NBH remains a globally significant marine fishing port.^1^ The uncontrolled discharge of polychlorinated biphenyls (PCBs) from capacitor manufacturing operations into the harbor during the mid-1900s resulted in severe PCB contamination, which was first detected in the mid-1970s at concentrations as high as 100,000 milligrams per kilogram dry-weight (mg/kg dw)^2^ in sediments, 53 mg/kg wet-weight (ww) in shellfish, and 22 mg/kg ww in finfish fillets.^3^ The contamination led to the harbor’s designation as a federal Superfund Site in 1983. Early investigations revealed a gradient of decreasing sediment PCB concentrations with increased distance from the source and resulted in designation of three management areas (Areas). Area 1, contains the original “hot spot” (PCBs > 4,000 mg/kg dw) and has undergone dredging since the late-1980s to remove PCB-contaminated sediments; Area 2, delineated at its downstream, southern edge by a hurricane barrier, has undergone navigational dredging; and Area 3 extends south from the hurricane barrier, into Buzzards Bay (**Figure 1**).

**Figure 1:**
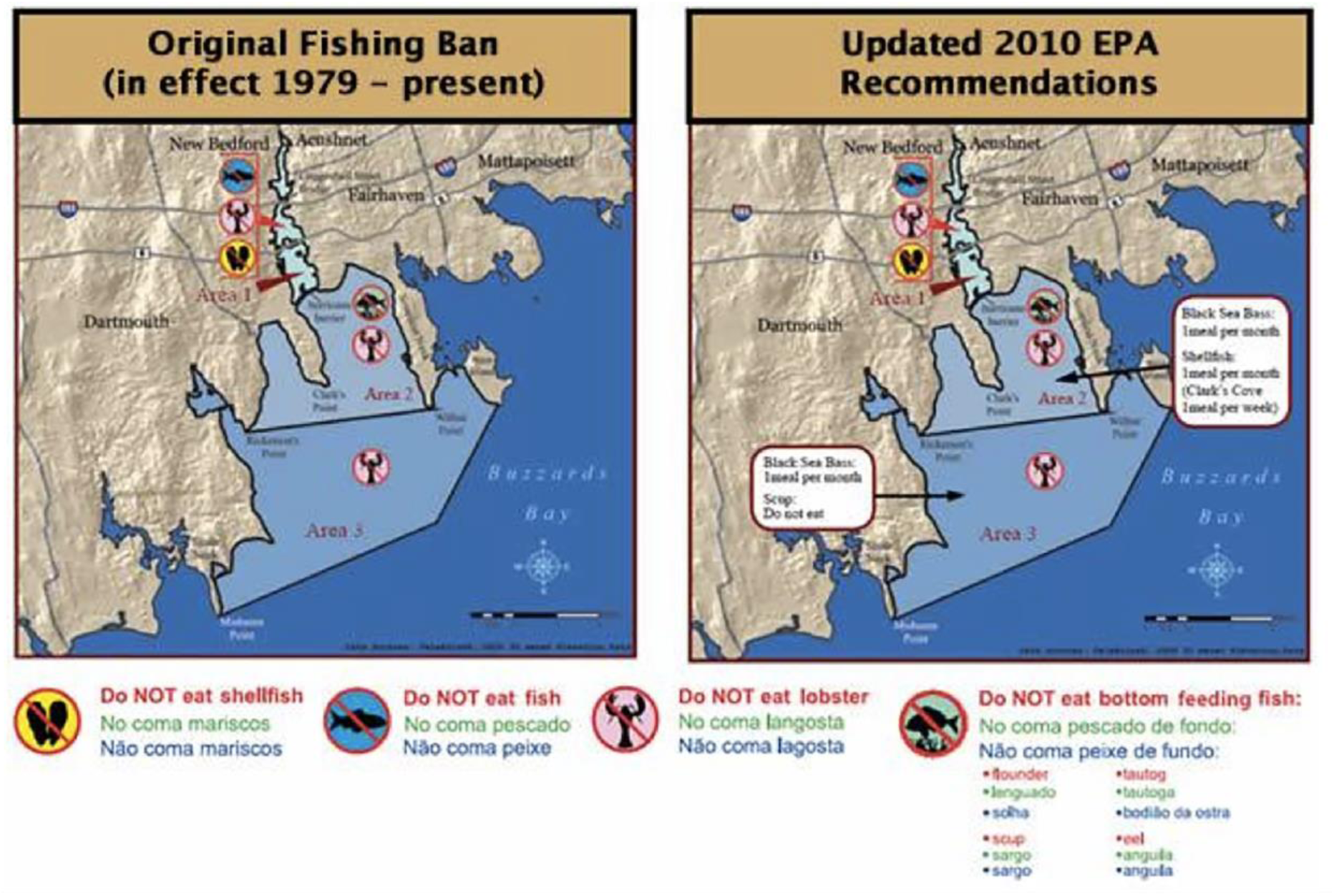
Map of New Bedford Harbor (NBH), MA, including delineations of the three seafood management Areas. Details of the NBH seafood advisory is available on the USEPA’s NBH Technical Documents website (https://www.epa.gov/new-bedford-harbor/fish-consumption-regulations-and-recommendations).

Consumption of seafood caught in Area 1 is prohibited and restrictions become less stringent in Areas 2 and 3 (**Figure 1**). Despite these restrictions, consumption of NBH-caught seafood continues^4^ amongst vulnerable populations living in surrounding communities, who may also be high-frequency fish consumers.^5^ The U.S. Environmental Protection Agency (USEPA) developed a site-specific threshold of 0.02 mg/kg ww for PCBs in consumable NBH seafood based on health risk analyses, accounting for high-frequency seafood consumption by the population around NBH.^6^ Because biota PCB concentrations are strongly influenced by sediment PCB concentrations,^7–9^ one goal of ongoing sediment dredging in NBH is to reduce the human health risk of PCB exposure from consumable seafood.^6^ Sediment cleanup goals established for NBH range between 1 and 50 mg/kg dw, depending on location within the harbor.^6^

To monitor remediation effectiveness, seafood species caught in NBH and consumed (e.g., quahog, or hard-shell clam, *Mercenaria mercenaria*, and scup, *Stenotomus chrysops*) are sampled annually by the MA Department of Environmental Protection (MassDEP) and Division of Marine Fisheries (MDMF).^10, 11^ Aroclors and 136 PCB congeners have been analyzed in the edible portion of seafood species since 2003 with the goals of characterizing temporal changes in PCB levels and evaluating differences in PCB concentrations between seafood in the three Areas. A number of studies have quantified temporal changes in PCB concentrations in wildlife,^12, 13^ but evaluating analogous trends in NBH is unique considering the harbor has undergone extensive remediation (projected to ultimately total 900,000 m^3^ of sediment). This study is the first to examine temporal and spatial trends of PCB concentrations in NBH quahogs and scup and to contextualize the trends using a health risk assessment to assess the progress of harbor clean-up efforts.

## 2. Methods

Detailed sampling procedures, analytical methods for lipid and PCB analysis, and data validation summaries are found in annual NBH seafood monitoring reports.^10, 11^

### 2.1 Seafood Collection and Data Sources

Quahogs and scup were collected by MDMF annually throughout the three NBH Areas since 2003 (**Figure 1**). All specimens were of legally harvestable size and collected during legal harvest seasons. Quahogs were collected pre-spawn to capture peak lipid concentration and to provide conservative estimates of PCBs in the edible tissue. Due to site conditions and spatial variability, organisms were not found nor sampled at every location each year. Collection and organism summary statistics are provided in **Table S1**.

We compiled historic data on PCB concentrations in seafood collected from 1976-1979.^3, 14, 15^ We aggregated data for quahogs with four shellfish species and for scup with seven bottom-feeding finfish species (**Table S2**). Shellfish are stationary and in intimate contact with sediment. Therefore, shellfish may closely reflect sediment PCBs over a small spatial scale. Finfish are mobile over a larger spatial area, in some instances migrating seasonally between coastal and off-shore waters. Finally, physiological differences between these groups might influence how they process PCBs to which they are exposed.^16^

### 2.2 PCB Analysis

Quahog composite samples consisted of approximately 12 individual whole, shelled organisms per location. Scup composites were comprised of edible, skin-on fillets from roughly five individual fish per location. Composite samples were extracted by USEPA Method 3570. Extracts were analyzed for lipid content, Aroclors and 136 PCB congeners based on USEPA Methods 680 and 8270D. Included in this suite of congeners are all non-*ortho* and mono-*ortho* substituted PCBs. Lipid content was reported as a weight-based percentage. PCB analytes were reported as mg/kg ww. Analytes detected outside the analytical calibration limits were reported as estimated (“J”) values. Analytes not detected above the sample quantitation limit (SQL) were reported as one half the SQL (half-SQL).

### 2.3 Data Analyses

Wet-weight PCB concentrations of individual congeners (PCB_WW_) in seafood were lipid-normalized (PCB_LN_). Total PCBs were calculated as the sum of individual congeners, including J-values and half-SQL, for both wet-weight (ΣPCB_WW_) and lipid-normalized (ΣPCB_LN_) concentrations. Individual PCB_LN_ congeners were grouped by homolog to evaluate congener patterns. The SQL for 73 congeners measured in quahogs decreased significantly (alpha=0.05) from 2003-2016, whereas only seven congener-specific SQLs declined in scup from 2003-2014. Although changes in SQL over time may influence temporal trends in PCBs levels, particularly in less contaminated regions of the harbor, half-SQL values were included to provide conservative estimates of PCB concentrations.

### 2.4 Health Risk Assessment

We evaluated the health risk of consuming NBH-caught seafood for two time periods: Past, 1980, prior to the harbor’s Superfund designation; and Present, the three most recent years each species was sampled by MassDEP (“present”) and 2015. Exposure Point Concentrations (EPCs), the PCB concentrations in seafood that people are expected to consume, were calculated for ΣPCB_WW_, consistent with PCB EPCs used in USEPA risk calculations. EPCs were calculated for quahogs and scup at each time point, in each Area. EPCs for 1980 and 2015were estimated by linear regression equations describing the temporal change in log-transformed ΣPCB_WW_ (quahogs: 2003-2016; scup: 2003-2014). “Present” EPCs were calculated using the most recent empirical ΣPCB_WW_ data for each species (quahogs:2014-2016; scup: 2012-2014), selecting the 95% upper confidence limit (UCL) on the mean.^17^ Using USEPA exposure and toxicity assumptions from the NBH seafood advisory (**Table S3**),^18^ we calculated excess lifetime cancer risk (ELCR) from consuming one (Central Tendency Exposure, CTE) or four (Reasonable Maximum Exposure, RME) meals of NBH quahogs and scup per month.

### 2.5 Statistical Analyses

Statistical analyses were performed using R, version 3.6.0.^19^ ANOVA and t-tests were used to evaluate differences in mean seafood PCB levels between Areas. Linear regression was used to evaluate changes in PCB concentrations over time. ΣPCB_WW_ were log-transformed to meet the normality assumption of linear regression and prevent bias in interpreting linear regression parameters. Statistical significance was evaluated using alpha=0.05.

## 3. Experimental Results and Discussion

### 3.1 PCB congener detection, concentrations and patterns

Summary statistics show that ΣPCB_LN_ in the three Areas ranged from 7.55-1,480 mg/kg in quahogs (2003-2016) and from 2.96-250 mg/kg in scup (2003-2014) (**Table S1**). The frequency of congener detection above the SQL ranged from 0-97% in quahogs and 0-100% in scup, with 16 and 48 congeners detected in at least 90% of quahog and scup samples, respectively. Three congeners (PCBs 76, 184, 205) were not detected in any sample for either species. Dioxin-like PCBs comprise up to 10.8% of ΣPCB_LN_ in quahogs and up to 20.3% of ΣPCB_LN_ in scup (**Figure S1**).

The distributions of PCB_LN_ of congeners within each homolog show that quahogs are dominated by tri- to hexa-chlorobiphenyls (CBs), with the highest mean congener concentration measured in tri-CBs (**Figure 2a**). In contrast, PCBs in scup are dominated by tetra- to hexa-CBs, with highest mean congener concentrations in penta- and hexa-CBs (**Figure 2b**). NBH PCB contamination resulted primarily from release of Aroclors-1242 and −1016, and to a lesser extent −1254.^3^ The homolog pattern in quahogs from Areas 1 and 2 most closely resembles weathered Aroclors-1242 and −1016 (**Table S4**), characterized by the disproportionate loss of lighter congeners over time. Since bivalves are stationary and have negligible capacity for xenobiotic biotransformation,^20^ the PCB congener pattern found in quahogs is considered a proxy for *in-situ* conditions.

**Figure 2:**
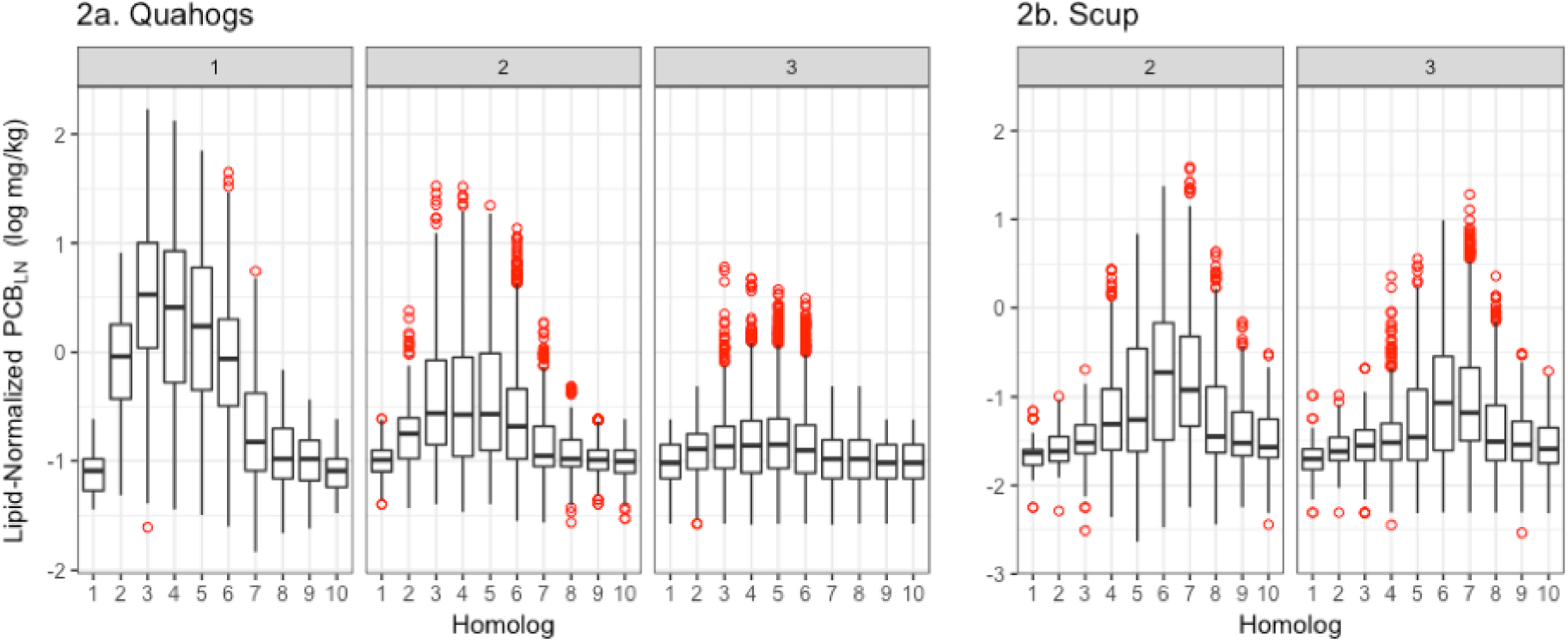
Distribution of ΣPCB_LN_ (log mg/kg ww) concentrations measured in quahogs (a) and scup (b) grouped by number of chlorines (homolog) for all years in each Area. Data are reported as mean ± interquartile range (IQR) of individual congeners within each homolog for 38-80 samples collected in a given area between 2003-2016 for quahogs and 2003-2014 for scup. Outliers are depicted.

The abundance of penta- and hexa-CBs in NBH scup (**Table S4**) can either be explained by Aroclor-1254 discharge into NBH^3^ or the tendency for highly chlorinated congeners to bioaccumulate in aquatic environments,^16^ biomagnifying in higher trophic organisms,^7–9^ like scup. The presence of hydroxylated PCB metabolites in scup^21^ suggests that scup physiology also may contribute to the relative absence of di-, tri- and tetra-CBs since less chlorinated congeners are generally most amenable to biotransformation. Overall, the difference in homolog signatures between quahogs and scup likely reflects their respective ecological niches, trophic positions, and biotransformation capacity.

### 3.3 Spatial and temporal patterns of PCBs in NBH seafood

Spatially, ΣPCB_LN_ in quahogs decreases with increased distance from the PCB source (Area 1 > Area 2 > Area 3, ANOVA p≤0.0001) when data are pooled by Area across all years (**Figure 3a**). The same pattern was observed in scup (**Figure 3b**; Area 2 > Area 3, T-test p≤0.0001). Trends for dioxin-like and non-dioxin-like ΣPCB_LN_ were similar (**Figure S1**). These findings are consistent with spatial PCB gradients previously demonstrated in other NBH shellfish^3^ and finfish.^3, 22^

**Figure 3:**
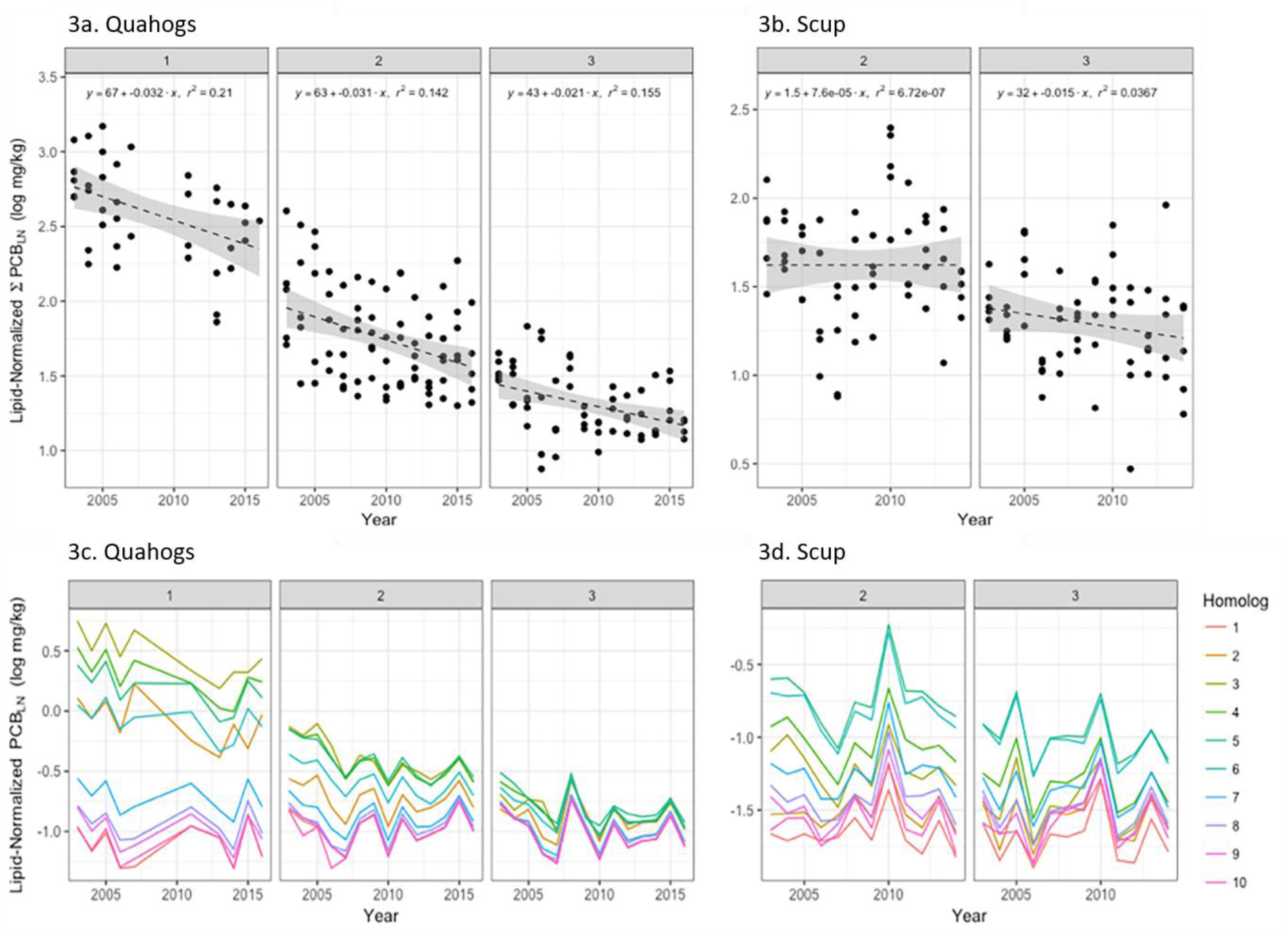
Linear regression analysis of changes in ΣPCB_LN_ (log mg/kg ww) over time in each NBH seafood management Area in quahogs (a) and scup (b). N = 1 to 7 composite samples/Area/year for quahogs and n = 5 composite samples/Area/year for scup. PCB_LN_ (c, d) show the annual mean concentration of PCB congeners within each homolog in quahogs and scup in a given year and Area. Included in ΣPCB_LN_ are all non-*ortho* (PCBs 77, 81, 126, 169) and mono-*ortho* (PCBs 105, 114, 118, 123, 156, 157, 167, 189) substituted PCBs, as well as all National Oceanographic and Atmospheric Administration (NOAA) and World Health Organization (WHO) congeners. Complete homolog data is presented in Table S5.

Concentrations of ΣPCB_LN_ were log-transformed in order to evaluate temporal trends in quahogs (2003-2016) and scup (2003-2014) in each Area using linear regression. Temporally, ΣPCB_LN_ in quahogs declined significantly (Area 1: −3.2% mg/kg/year, p=0.0039; Area 2: −3.1% mg/kg/year, p=0.0006; Area 3: −2.1% mg/kg/year, p=0.0017) (**Figure 3a**). However, significant declines in ΣPCB_LN_ were not observed in scup (**Figure 3b**). PCB concentrations in stationary shellfish reflect anticipated changes in *in-situ* PCB levels over time, following extensive sediment remediation, whereas mobile finfish do not.

Temporal declines in PCB_LN_ for individual congeners in quahogs are significant amongst dominant homolog groups (**Figure 3c, Table S5**). The absence of significant declines in other homologs is likely attributable to lower congener detection frequencies and a higher prevalence of non-detect congeners substituted with half-SQL. PCB_LN_ in scup declined significantly in tetra-CBs in Area 2 and tetra- to septa-CBs in Area 3 (**Figure 3d, Table S5**). Together, these data show that PCB concentrations in NBH seafood decrease with time and distance from the PCB source. Declining trends in PCB concentrations observed in NBH seafood since 2003, particularly quahogs in Area 1, suggest that sediment remediation is associated with improvements in environmental quality. Declining PCB concentrations may also be attributed to natural PCB “weathering” processes such as degradation, volatilization and settling into sediments.

Differences in sample collection, handling and analysis preclude quantitative comparison of historic (1970s) and current NBH seafood PCB concentrations, however, doing so qualitatively is useful to provide context for PCB body burdens in edible seafood at the peak of PCB contamination in NBH. To this end, we aggregated available, historic PCB data from NBH shellfish and bottom-dwelling finfish collected between 1976-1979 (**Table S2)**. PCB concentrations in NBH seafood were reported as Aroclor-1254 (mg/kg ww). The highest concentration of Aroclor-1254 measured in quahogs in the late-1970s was 4.0 mg/kg in Area 1;^3^ whereas concentrations as high as 53 mg/kg were reported in soft-shell clams collected in Area 1.^14^ The spatial gradient in PCB concentration reported above also was evident among NBH shellfish collected during the late-1970s (mean Aroclor-1254 in Area 1 > Area 2 > Area 3). The highest Aroclor-1254 concentration reported in scup was 11.4 mg/kg in Area 2.^3, 14^ In bottom-dwelling finfish, the highest Aroclor-1254 concentration was 22 mg/kg in winter flounder in Area 1.^3^ While PCB concentrations in scup did not follow the spatial gradient observed with shellfish, PCB concentrations in aggregated bottom-dwelling finfish were consistent (mean Aroclor-1254 in Area 1 > Area 2 > Area 3). In comparison, the maximum Aroclor-1254 measured between 2003-2013 was 2.40 mg/kg in quahogs in Area 1 and 3.65 mg/kg in scup in Area 2.

### 3.4 Comparative risk assessment for fish consumption over time

To analyze the human health impact of decreasing PCB concentrations in NBH seafood, we calculated historic and current cancer risk from consuming NBH-caught seafood. First, we determined the 95% UCL of the mean ΣPCB_WW_ concentration within the most recent three years of data (quahogs: 2014-2016, scup: 2012-2014) in each Area to serve as EPCs for “present” risk calculations. Second, we used linear regression analysis to evaluate changes in ΣPCB_WW_ measured in NBH quahogs and scup since 2003. The resulting regression parameters were then used to estimate PCB concentrations in NBH seafood in 1980 and 2015 (**Table S6** and **Figure S2)** for all Areas, which served as EPCs. We calculated the cancer risk associated with consuming one (CTE) or four (RME) meals of quahogs or scup per month (**Table 1**) relative to the risk criterion of 1 E-4 used by USEPA for NBH. Temporal and spatial trends in cancer risk for quahogs and scup mimic the directionality of ΣPCB_WW_ changes over time due to the linearity of models used to derive EPCs and cancer risk. For example, we show that in Area 2, where ΣPCB_WW_ has been declining at a rate of −3.4% mg/kg/yr (2003-2016) in quahogs, the CTE cancer risk level for adults declined from 2.16 E-4 in 1980, to 1.36 E-5 in 2015 (**Table 1**). The “present” cancer risk for adults consuming quahogs harvested in Area 2 is 2.31 E-5. By contrast, ΣPCB_WW_ in Area 2 scup increased by 0.28% mg/kg/yr (2003-2014), resulting in CTE cancer risk increases for adults from a risk level of 6.38 E-5 in 1980, to risk levels of 7.97 E-5 in 2015 (**Table 1**). The “present” cancer risk for adults consuming scup harvested in Area 2 is 1.66 E-4.

**Table 1:** Summary of human cancer risk for PCB exposures associated with the consumption of quahogs and scup (b) harvested from NBH for young children (1-6 years) and adults (16-70 years). Cancer risk was calculated using USEPA exposure and toxicity assumptions in Table S3, the Exposure Point Concentrations (EPCs) for ΣPCB_WW_ (mg/kg ww) shown in Table S6, and the oral cancer slope factor for total PCBs (2.0 (mg/kg/day)^−1^).^24^ Cancer risk was calculated for consumption of one (Central Tendency Exposure, CTE) or four (Reasonable Maximum Exposure, RME) meals of NBH quahogs and scup per month. All cancer risk estimates were evaluated relative to the health risk criterion of 1 E-4 (indicated by the red line) used by USEPA for NBH.

| Species | Date | Exposure Scenario | Area 1 |  |  |  | Area 2 |  |  |  | Area 3 |  |  |  |
| --- | --- | --- | --- | --- | --- | --- | --- | --- | --- | --- | --- | --- | --- | --- |
|  |  |  | Young Child |  | Adult |  | Young Child |  | Adult |  | Young Child |  | Adult |  |
|  |  |  | LADD | ELCR | LADD | ELCR | LADD | ELCR | LADD | ELCR | LADD | ELCR | LADD | ELCR |
| Quahogs | Present (2014-2016) | CTE | 2.55E-05 | 5.10E-05 | 1.20E-04 | 2.39E-04 | 2.46E-06 | 4.93E-06 | 1.16E-05 | 2.31E-05 | 8.33E-07 | 1.67E-06 | 3.91E-06 | 7.82E-06 |
|  |  | RME | 1.11E-04 | 2.21E-04 | 5.19E-04 | 1.04E-03 | 1.07E-05 | 2.13E-05 | 5.01E-05 | 1.00E-04 | 3.61E-06 | 7.22E-06 | 1.70E-05 | 3.39E-05 |
|  | 1980 | CTE | 2.05E-04 | 4.09E-04 | 9.60E-04 | 1.92E-03 | 2.31E-05 | 4.61E-05 | 1.08E-04 | 2.16E-04 | 1.89E-06 | 3.79E-06 | 8.89E-06 | 1.78E-05 |
|  |  | RME | 8.86E-04 | 1.77E-03 | 4.16E-03 | 8.32E-03 | 9.99E-05 | 2.00E-04 | 4.69E-04 | 9.38E-04 | 8.21E-06 | 1.64E-05 | 3.85E-05 | 7.71E-05 |
|  | 2015 | CTE | 1.20E-05 | 2.39E-05 | 5.62E-05 | 1.12E-04 | 1.45E-06 | 2.90E-06 | 6.81E-06 | 1.36E-05 | 7.50E-07 | 1.50E-06 | 3.52E-06 | 7.04E-06 |
|  |  | RME | 5.19E-05 | 1.04E-04 | 2.43E-04 | 4.87E-04 | 6.29E-06 | 1.26E-05 | 2.95E-05 | 5.90E-05 | 3.25E-06 | 6.50E-06 | 1.52E-05 | 3.05E-05 |
| Scup | Present (2012-2014) | CTE | -- | -- | -- | -- | 1.77E-05 | 3.54E-05 | 8.30E-05 | 1.66E-04 | 7.16E-06 | 1.43E-05 | 3.36E-05 | 6.72E-05 |
|  |  | RME | -- | -- | -- | -- | 7.66E-05 | 1.53E-04 | 3.60E-04 | 7.19E-04 | 3.10E-05 | 6.20E-05 | 1.46E-04 | 2.91E-04 |
|  | 1980 | CTE | -- | -- | -- | -- | 6.80E-06 | 1.36E-05 | 3.19E-05 | 6.38E-05 | 1.21E-05 | 2.41E-05 | 5.66E-05 | 1.13E-04 |
|  |  | RME | -- | -- | -- | -- | 2.95E-05 | 5.89E-05 | 1.38E-04 | 2.77E-04 | 5.23E-05 | 1.05E-04 | 2.45E-04 | 4.91E-04 |
|  | 2015 | CTE | -- | -- | -- | -- | 8.49E-06 | 1.70E-05 | 3.99E-05 | 7.97E-05 | 3.09E-06 | 6.18E-06 | 1.45E-05 | 2.90E-05 |
|  |  | RME | -- | -- | -- | -- | 3.68E-05 | 7.36E-05 | 1.73E-04 | 3.45E-04 | 1.34E-05 | 2.68E-05 | 6.29E-05 | 1.26E-04 |

We acknowledge several limitations of our evaluation of temporal changes in human health risk from PCB exposure through NBH-harvested seafood. Notably, USEPA calculated ΣPCB_WW_ substituting “zero” concentration for non-detect congeners, whereas we substituted half-SQL. USEPA also calculated EPCs using ΣPCB_WW_ data for quahogs and scup measured over a longer period of time. Despite differences in EPCs, our 2015 and “present” risk calculations in both quahogs and scup are congruent with existing Massachusetts Department of Public Health and USEPA fish consumption advisories for all Areas^23^ (**Figure 1**). Additionally, risk estimates for 1980 rely on the assumption that trends in ΣPCB_WW_ observed in quahogs and scup since 2003 can be generalized to describe the pattern of change in PCB concentrations over the past 40 years. Finally, we expect the rate of PCB removal from NBH during periods of active sediment remediation will differ from that of natural attenuation. Trends in ΣPCB_WW_ from 2003-2016 reflect changes in ΣPCB_WW_ during periods of active sediment remediation and may overestimate the rate of decline which has occurred since 1980 since remediation has not been steadily ongoing throughout this entire time period. However, it is important to note that significant sediment remediation operations did occur prior to 2003, including dredging of the most heavily contaminated sediments in the “hot spot” during 1994-1995, so we believe that changes ΣPCB_WW_ between 2003 and 2016 offer a relevant, though imprecise, estimate of trends predating the time period for which empirical data exists.

In summary, we show significant declines in ΣPCB_LN_ concentrations in quahogs over time and with increasing distance from the PCB source in Area 1. Temporal declines in PCB concentrations have led to a reduction in human cancer risk from consumption of NBH quahogs between 1980 and the present. Temporal trends in PCB concentrations in scup have changed less over time, likely due to their life cycle characteristics and mobility. By extension, scup offer less clarity about temporal changes in PCB concentrations throughout NBH and the associated cancer risk posed to humans by consuming NHB-harvested scup. Quahogs are expected to more accurately reflect changes in the environmental quality because they are sessile and of lower trophic status. Decreases in quahog PCB concentrations reflect extensive sediment dredging by the USEPA to reduce the total mass of PCB in NBH. We recommend that annual PCB congener and Aroclor monitoring be continued in quahogs all allow for future analysis of how PCB levels are changing in NBH seafood. We also recommend that annual PCB monitoring should be resumed in other species, such as scup, since high annual variability of PCB concentrations within mobile finfish necessitates robust datasets to accurately evaluate effects of ongoing sediment dredging on PCB levels in higher trophic organisms. Comprehensive evaluation of remediation efficacy also should include NBH-wide PCB fate and transport modeling, integrating available PCB sediment and air data with seafood PCB data.

## 5. Acknowledgments

KC, JS and WHB were funded by the Boston University Superfund Research Program, National Institute of Environmental Health Sciences Grant P42ES007381. The authors thank D. Lederer at USEPA Region 1, and B. Bergen and D. Nacci at USEPA Office of Research and Development, Atlantic Ecology Division for their review of the manuscript and for sharing technical details about the site history and ongoing PCB remediation.

