## Supplemental Materials for "Spatiotemporal trends in polychlorinated biphenyl concentrations in seafood based on long-term monitoring and remediation in New Bedford Harbor, Massachusetts"

### Table of Contents

**Page 2: Table S1:** Summary statistics for quahogs and scup describing sampling frequency, composite samples, lipid content, $\sum PCB_{WW}$ and $\sum PCB_{LN}$ by NBH seafood management Area and year.

**Page 3: Table S2.** Historic concentrations of Aroclor 1254 (mg/kg ww) in shellfish and bottom-dwelling finfish harvested from NBH between 1976 and 1979.

**Page 4: Table S3.** Exposure and toxicity assumptions used by EPA to evaluate human cancer risk from consuming NBH-harvested seafood.

**Page 5: Table S4.** Homologs as a percent of $\sum PCB_{LN}$ by weight in NBH quahogs and scup, and Aroclors discharged to NBH.

**Page 6: Table S5.** Linear regression parameters describing temporal changes in mean $PCB_{LN}$ (mg/kg ww) by homolog group in each Area for quahogs (2003-2016) and scup (2003-2014).

**Page 7: Table S6:** Linear regression parameters and Exposure Point Concentrations (EPCs) for $\sum PCB_{WW}$ (mg/kg ww) in NBH quahogs and scup.

**Page 8: Figure S1:** Concentrations in dioxin-like and non-dioxin like $\sum PCB_{LN}$ (log mg/kg ww) in quahogs (a) and scup (b) in each seafood management Area over time.

**Page 9: Figure S2.** Linear regressions for changes in $\sum PCB_{WW}$ (log mg/kg ww) over time used to estimate PCB concentrations in NBH quahogs (a) and scup (b) in 1980 and 2015.

**Table S1:** Summary statistics for quahogs and scup describing sampling frequency, composite samples, lipid content, $\sum PCB_{WW}$ and $\sum PCB_{LN}$ by NBH seafood management Area and year.

| Species | Area | Year | 2003 | 2004 | 2005 | 2006 | 2007 | 2008 | 2009 |
|---|---|---|---|---|---|---|---|---|---|
| Quahog | 1 | No. Locations Sampled (Sampled/Total) | 5/5 | 5/5 | 5/5 | 5/5 | 2/5 | 0/5 | 0/5 |
| (*Mercenaria* | | No. Individuals per Composite Sample (Mean (Min, Max)) | 14 (13,15) | 15 (13,19) | 12 (12,12) | 12 (12,12) | 12 (12,12) | na | na |
| *mercenaria*) | | Body Lipids, % (GM (Mean, SD)) | 0.21 (0.22, 0.08) | 0.37 (0.37, 0.08) | 0.24 (0.24, 0.04) | 0.46 (0.47, 0.10) | 0.51 (0.51, 0.07) | na | na |
| | | $\sum PCB_{WW}$, mg/kg ww (GM (Mean, SD)) | 1.42 (1.58, 0.82) | 1.60 (2.28, 2.25) | 1.61 (1.88, 1.25) | 1.61 (1.93, 1.32) | 2.75 (3.64, 3.38) | na | na |
| | | $\sum PCB_{LN}$, mg/kg ww (GM (Mean, SD)) | 676 (716, 289) | 439 (564, 441) | 667 (778, 474) | 351 (409, 259) | 541 (675, 570) | na | na |
| | 2 | No. Locations Sampled (Sampled/Total) | 5/9 | 5/9 | 5/9 | 5/9 | 6/9 | 6/9 | 6/9 |
| | | No. Individuals per Composite Sample (Mean (Min, Max)) | 13 (12,14) | 13 (7,15) | 12 (12,12) | 12 (12,12) | 12 (12,12) | 12 (12,12) | 12 (12,12) |
| | | Body Lipids, % (GM (Mean, SD)) | 0.15 (0.16, 0.06) | 0.20 (0.23, 0.13) | 0.21 (0.22, 0.05) | 0.30 (0.31, 0.09) | 0.39 (0.40, 0.11) | 0.20 (0.21, 0.06) | 0.16 (0.17, 0.05) |
| | | $\sum PCB_{WW}$, mg/kg ww (GM (Mean, SD)) | 0.17 (0.28, 0.35) | 0.19 (0.28, 0.29) | 0.22 (0.34, 0.31) | 0.22 (0.28, 0.20) | 0.17 (0.24, 0.25) | 0.12 (0.14, 0.11) | 0.10 (0.12, 0.10) |
| | | $\sum PCB_{LN}$, mg/kg ww (GM (Mean, SD)) | 113 (152, 145) | 97 (136, 120) | 103 (149, 116) | 73 (85, 51) | 45 (54, 39) | 59 (71, 45) | 60 (67, 37) |
| | 3 | No. Locations Sampled (Sampled/Total) | 5/9 | 5/9 | 5/9 | 5/9 | 5/9 | 4/9 | 4/9 |
| | | No. Individuals per Composite Sample (Mean (Min, Max)) | 15 (12,19) | 17 (13,20) | 12 (12,12) | 12 (12,12) | 12 (12,12) | 12 (12,12) | 12 (12,12) |
| | | Body Lipids, % (GM (Mean, SD)) | 0.14 (0.15, 0.06) | 0.19 (0.19, 0.05) | 0.21 (0.22, 0.05) | 0.34 (0.34, 0.05) | 0.42 (0.45, 0.21) | 0.12 (0.13, 0.04) | 0.22 (0.22, 0.03) |
| | | $\sum PCB_{WW}$, mg/kg ww (GM (Mean, SD)) | 0.05 (0.05, 0.02) | 0.05 (0.06, 0.01) | 0.05 (0.06, 0.05) | 0.08 (0.11, 0.09) | 0.06 (0.07, 0.03) | 0.05 (0.05, 0) | 0.04 (0.04, 0) |
| | | $\sum PCB_{LN}$, mg/kg ww (GM (Mean, SD)) | 35.12 (35.57, 6.46) | 28.64 (29.83, 9.13) | 24.77 (29.21, 21.89) | 22.46 (31.73, 26.08) | 14.79 (16, 7.76) | 36.47 (37.14, 7.77) | 16.34 (16.49, 2.59) |
| Scup | 2 | No. Locations Sampled (Sampled/Total) | 5/5 | 5/5 | 5/5 | 5/5 | 5/5 | 5/5 | 5/5 |
| (*Stenotomus* | | No. Individuals per Composite Sample (Mean (Min, Max)) | 3 (3,3) | 5 (5,5) | 5 (5,5) | 5 (5,5) | 5 (4,5) | 5 (5,5) | 5 (5,5) |
| *chrysops*) | | Muscle Lipids, % (GM (Mean, SD)) | 1.09 (1.11, 0.28) | 1.2 (1.35, 0.67) | 1.11 (1.15, 0.3) | 1.29 (1.57, 1.31) | 1.13 (1.19, 0.46) | 0.81 (0.82, 0.2) | 1.16 (1.17, 0.13) |
| | | $\sum PCB_{WW}$, mg/kg ww (GM (Mean, SD)) | 0.67 (0.78, 0.4) | 0.66 (0.8, 0.55) | 0.48 (0.55, 0.31) | 0.33 (0.39, 0.24) | 0.18 (0.21, 0.13) | 0.28 (0.34, 0.25) | 0.4 (0.45, 0.23) |
| | | $\sum PCB_{LN}$, mg/kg ww (GM (Mean, SD)) | 62.16 (70.11, 37.29) | 55.3 (57.9, 19.94) | 43.38 (46.93, 19.56) | 25.23 (33.55, 27.88) | 15.64 (18.58, 11.15) | 34.74 (41.97, 28.29) | 34.59 (37.68, 16.36) |
| | 3 | No. Locations Sampled (Sampled/Total) | 5/5 | 5/5 | 5/5 | 5/5 | 5/5 | 5/5 | 5/5 |
| | | No. Individuals per Composite Sample (Mean (Min, Max)) | 3 (3,3) | 5 (5,5) | 5 (5,5) | 5 (5,5) | 5 (5,5) | 5 (5,5) | 5 (5,5) |
| | | Muscle Lipids, % (GM (Mean, SD)) | 0.91 (0.92, 0.14) | 1.64 (1.64, 0.09) | 1.07 (1.15, 0.55) | 1.82 (1.92, 0.74) | 1 (1.02, 0.21) | 1.1 (1.13, 0.24) | 0.98 (1.05, 0.41) |
| | | $\sum PCB_{WW}$, mg/kg ww (GM (Mean, SD)) | 0.24 (0.26, 0.11) | 0.31 (0.32, 0.05) | 0.45 (0.57, 0.46) | 0.19 (0.2, 0.08) | 0.19 (0.22, 0.12) | 0.21 (0.22, 0.08) | 0.19 (0.2, 0.07) |
| | | $\sum PCB_{LN}$, mg/kg ww (GM (Mean, SD)) | 26.6 (27.52, 8.68) | 19.07 (19.33, 3.62) | 42.07 (45.99, 19.28) | 10.4 (10.55, 1.85) | 19.16 (21.35, 11.2) | 19.26 (19.77, 4.88) | 18.98 (22.27, 12.03) |

| Species | Area | Year | 2010 | 2011 | 2012 | 2013 | 2014 | 2015 | 2016 |
|---|---|---|---|---|---|---|---|---|---|
| Quahog | 1 | No. Locations Sampled (Sampled/Total) | 0/5 | 4/5 | 0/5 | 5/5 | 5/5 | 3/5 | 1/5 |
| (*Mercenaria* | | No. Individuals per Composite Sample (Mean (Min, Max)) | na | 14 (13,14) | na | 13 (13,13) | 13 (13,13) | 12 (12,12) | 12 |
| *mercenaria*) | | Body Lipids, % (GM (Mean, SD)) | na | 0.20 (0.2, 0.05) | na | 0.27 (0.30, 0.13) | 0.49 (0.49, 0.02) | 0.16 (0.17, 0.07) | 0.38 |
| | | $\sum PCB_{WW}$, mg/kg ww (GM (Mean, SD)) | na | 0.72 (0.80, 0.41) | na | 0.52 (0.59, 0.31) | 1.25 (1.35, 0.65) | 0.55 (0.57, 0.20) | 1.31 |
| | | $\sum PCB_{LN}$, mg/kg ww (GM (Mean, SD)) | na | 359 (412, 238) | na | 189 (269, 233) | 256 (279, 147) | 333 (341, 90) | 345 |
| | 2 | No. Locations Sampled (Sampled/Total) | 6/9 | 7/9 | 6/9 | 9/9 | 9/9 | 6/9 | 6/9 |
| | | No. Individuals per Composite Sample (Mean (Min, Max)) | 12 (12,13) | 13 (13,16) | 12 (12,13) | 13 (13,13) | 13 (13,13) | 12 (12,12) | 12 (12,12) |
| | | Body Lipids, % (GM (Mean, SD)) | 0.36 (0.37, 0.09) | 0.18 (0.19, 0.06) | 0.26 (0.27, 0.07) | 0.26 (0.26, 0.04) | 0.22 (0.23, 0.07) | 0.13 (0.13, 0.05) | 0.22 (0.22, 0.02) |
| | | $\sum PCB_{WW}$, mg/kg ww (GM (Mean, SD)) | 0.14 (0.18, 0.14) | 0.10 (0.13, 0.12) | 0.12 (0.14, 0.11) | 0.09 (0.11, 0.08) | 0.10 (0.12, 0.10) | 0.07 (0.08, 0.05) | 0.08 (0.10, 0.07) |
| | | $\sum PCB_{LN}$, mg/kg ww (GM (Mean, SD)) | 39 (48, 38) | 57 (74, 57) | 45 (50, 29) | 35 (391, 24) | 45 (53, 38) | 58 (74, 60) | 38 (44, 31) |
| | 3 | No. Locations Sampled (Sampled/Total) | 4/9 | 4/9 | 4/9 | 9/9 | 9/9 | 4/9 | 4/9 |
| | | No. Individuals per Composite Sample (Mean (Min, Max)) | 13 (12,13) | 13 (12,14) | 14 (9,23) | 13 (13,13) | 13 (13,13) | 12 (12,12) | 12 (12,12) |
| | | Body Lipids, % (GM (Mean, SD)) | 0.38 (0.41, 0.17) | 0.2 (0.21, 0.09) | 0.3 (0.31, 0.09) | 0.29 (0.31, 0.11) | 0.26 (0.28, 0.1) | 0.17 (0.20, 0.12) | 0.30 (0.3, 0.06) |
| | | $\sum PCB_{WW}$, mg/kg ww (GM (Mean, SD)) | 0.05 (0.05, 0.01) | 0.04 (0.04, 0.01) | 0.05 (0.05, 0.01) | 0.05 (0.05, 0.01) | 0.04 (0.04, 0.01) | 0.04 (0.04, 0.01) | 0.04 (0.04, 0.01) |
| | | $\sum PCB_{LN}$, mg/kg ww (GM (Mean, SD)) | 13.26 (13.47, 2.67) | 19.72 (20.33, 5.59) | 16.96 (17.35, 4.36) | 16.07 (16.86, 6.23) | 16.56 (17.98, 9.37) | 23.31 (24.47, 8.67) | 14.18 (14.29, 1.98) |
| Scup | 2 | No. Locations Sampled (Sampled/Total) | 5/5 | 5/5 | 5/5 | 5/5 | 5/5 | ns | ns |
| (*Stenotomus* | | No. Individuals per Composite Sample (Mean (Min, Max)) | 5 (5,5) | 5 (4,5) | 5 (5,5) | 5 (5,5) | 5 (5,5) | ns | ns |
| *chrysops*) | | Muscle Lipids, % (GM (Mean, SD)) | 0.53 (0.58, 0.3) | 1.14 (1.16, 0.24) | 1.67 (1.8, 0.77) | 0.9 (0.91, 0.14) | 1.57 (1.59, 0.24) | ns | ns |
| | | $\sum PCB_{WW}$, mg/kg ww (GM (Mean, SD)) | 0.77 (0.83, 0.35) | 0.54 (0.62, 0.38) | 0.82 (1.06, 0.8) | 0.36 (0.43, 0.26) | 0.49 (0.51, 0.18) | ns | ns |
| | | $\sum PCB_{LN}$, mg/kg ww (GM (Mean, SD)) | 145.57 (163.36, 76.83) | 47.34 (56.1, 39.8) | 49.25 (53.69, 22.88) | 39.64 (48.45, 29.24) | 30.94 (31.71, 7.48) | ns | ns |
| | 3 | No. Locations Sampled (Sampled/Total) | 5/5 | 5/5 | 5/5 | 5/5 | 5/5 | ns | ns |
| | | No. Individuals per Composite Sample (Mean (Min, Max)) | 5 (5,5) | 5 (5,6) | 5 (5,6) | 5 (4,5) | 5 (4,5) | ns | ns |
| | | Muscle Lipids, % (GM (Mean, SD)) | 0.46 (0.49, 0.18) | 1.58 (1.88, 1.45) | 1.61 (1.64, 0.38) | 0.87 (0.89, 0.24) | 1.42 (1.45, 0.33) | ns | ns |
| | | $\sum PCB_{WW}$, mg/kg ww (GM (Mean, SD)) | 0.17 (0.18, 0.07) | 0.19 (0.23, 0.14) | 0.25 (0.28, 0.12) | 0.2 (0.33, 0.41) | 0.19 (0.21, 0.09) | ns | ns |
| | | $\sum PCB_{LN}$, mg/kg ww (GM (Mean, SD)) | 36.06 (39.57, 19.85) | 12.31 (16.37, 11.72) | 15.86 (17.01, 7.74) | 23.13 (32.53, 33.64) | 13.24 (15.33, 8.65) | ns | ns |

**Table S2.** Historic concentrations of Aroclor 1254 (mg/kg ww*) in shellfish and bottom-dwelling finfish harvested from NBH between 1976 and 1979 (data compiled from: Santos, 1978; Kolek & Ceurvels, 1981; and Nisbet & Reynolds, 1984). Instances where Statistic = "N/A" signify that only one Aroclor 1254 measurement was reported for a given species. *Note: Every effort was made to verify whether historic reports presented Aroclor 1254 as mg/kg ww or dw; data presented herein are assumed to be ww.

| **Shellfish, Bivalves** | | **Statistic** | **Area 1** | **Area 2** | **Area 3** | **Overall** |
|---|---|---|---|---|---|---|
| **Species** | | | | | | |
| Mussles | *Mytilus edulis* | N/A | 3.7 | -- | -- | -- |
| Oyster | *Crassostrea virginica* | N/A | 15.8 | -- | -- | -- |
| Quahog | *Mercinaria mercinaria* | Min | 1.6 | 0.2 | 0.3 | 0.2 |
| | | Mean | 2.6 | 1.0 | 0.4 | 1.1 |
| | | Median | 2.1 | 0.7 | 0.4 | 0.7 |
| | | Max | 4.0 | 3.3 | 0.6 | 4.0 |
| | | N | 3 | 12 | 5 | 20 |
| Soft-shelled clam | *Mya arenaria* | Min | 14.6 | -- | -- | 14.6 |
| | | Mean | 29.7 | -- | -- | 29.7 |
| | | Median | 22.0 | -- | -- | 22.0 |
| | | Max | 53.0 | -- | -- | 53.0 |
| | | N | 7 | -- | -- | 7 |
| Shellfish, Bivalves | All Species | Min | 1.6 | 0.2 | 0.3 | 0.2 |
| | | Mean | 21.0 | 1.0 | 0.4 | 8.6 |
| | | Median | 21.0 | 0.7 | 0.4 | 1.3 |
| | | Max | 53.0 | 3.3 | 0.6 | 53.0 |
| | | N | 11 | 12 | 5 | 29 |

| **Finfish, Bottom-dwelling** | | **Statistic** | **Area 1** | **Area 2** | **Area 3** | **Overall** |
|---|---|---|---|---|---|---|
| **Species** | | | | | | |
| Black sea bass | *Centropristis striata* | N/A | -- | -- | 0.4 | -- |
| Fourpsot flounder | *Hippoglossina oblonga* | N/A | -- | -- | 0.8 | -- |
| Scup | *Stenotomus chrysops* | Min | 2.3 | 6.1 | 0.0 | 0.0 |
| | | Mean | 4.2 | 9.6 | 0.4 | 4.8 |
| | | Median | 4.2 | 11.4 | 0.0 | 4.2 |
| | | Max | 6.1 | 11.4 | 1.3 | 11.4 |
| | | N | 2 | 3 | 3 | 8 |
| Summer flounder | *Paralichthys dentatus* | Min | 2.1 | 0.2 | 0.3 | 0.2 |
| | | Mean | 6.1 | 5.1 | 2.2 | 4.5 |
| | | Median | 6.1 | 7.1 | 1.2 | 4.0 |
| | | Max | 10.0 | 7.9 | 4.0 | 10.0 |
| | | N | 2 | 3 | 2 | 7 |
| Tautog | *Tautoga onitis* | Min | -- | 0.1 | 0.1 | 0.1 |
| | | Mean | -- | 1.3 | 0.7 | 1.0 |
| | | Median | -- | 1.2 | 0.7 | 0.9 |
| | | Max | -- | 4.6 | 1.1 | 4.6 |
| | | N | -- | 9 | 7 | 16 |
| Window pane flounder | *Scophthalmus aquosus* | Min | 5.4 | -- | 3.1 | 3.1 |
| | | Mean | 9.5 | -- | 3.1 | 6.9 |
| | | Median | 8.8 | -- | 3.1 | 5.4 |
| | | Max | 14.3 | -- | 3.1 | 22.0 |
| | | N | 3 | -- | 2 | 5 |
| Winter flounder | *Pseudopleuronectes americanus* | Min | 6.0 | 0.0 | 0.2 | 0.0 |
| | | Mean | 10.4 | 5.6 | 3.9 | 5.6 |
| | | Median | 8.1 | 5.6 | 1.0 | 3.9 |
| | | Max | 22.0 | 11.0 | 20.0 | 22.0 |
| | | N | 7 | 5 | 19 | 31 |
| Finfish, Bottom-dwelling | All Species | Min | 2.1 | 0.0 | 0.0 | 0.0 |
| | | Mean | 8.7 | 4.2 | 2.6 | 4.3 |
| | | Median | 7.9 | 2.6 | 0.9 | 1.6 |
| | | Max | 22.0 | 11.4 | 20.0 | 22.0 |
| | | N | 14 | 20 | 35 | 69 |

**Table S3.** Exposure and toxicity assumptions used by USEPA to evaluate human cancer risk from consuming NBH-harvested seafood (adapted from USEPA, 2015: Third Five-Year Review Report, Appendix D (Risk Assessment Updates)).

| Receptor | Age (yrs) | Exposure Condition | IR | FI | EF | ED | BW | AT-c | SF |
|---|---|---|---|---|---|---|---|---|---|
| **Adult** | 16-70 | CTE | 0.227 | 1 | 12 | 55 | 70 | 25550 | 2.0 |
| | | RME | 0.227 | 1 | 52 | 55 | 70 | 25550 | 2.0 |
| **Young Child** | 1-6 | CTE | 0.114 | 1 | 12 | 5 | 15 | 25550 | 2.0 |
| | | RME | 0.114 | 1 | 52 | 5 | 15 | 25550 | 2.0 |

CTE = Central Tendancy Exposure
RME = Reasonable Maximum Exposure
IR = Ingestion Rate (kg/meal)
FI = Fraction Ingestion from site (assumed to be 1, unitless)
ED = Exposure Duration (years)
EF = Exposure Frequency (meals/year)
BW = Body Weight (kg)
AT-c = Averaging Time, cancer (days, 70 years * 365 days/yr = 25,550 days)
SF= Oral cancer slope factor ($(mg/kg/day)^{-1}$ )

**Table S4.** Homologs as a percent of $\sum PCB_{LN}$ by weight in NBH quahogs and scup, and Aroclors (Frame et al., 1996) discharged to NBH. Homolog groups are denoted by number of chlorines (e.g Homolog 2 represents di-chlorobiphenyls). The contribution of each homolog to $\sum PCB_{LN}$ and Aroclor mixtures increases as cell shading changes from green to red. ND denotes homologs not detected in an Aroclor mixture.

| Homolog | Quahogs | | | Scup | | Aroclors | | |
|---|---|---|---|---|---|---|---|---|
| | Area 1 | Area 2 | Area 3 | Area 2 | Area 3 | 1016 | 1242 | 1254 |
| 1 | 0.1 | 0.5 | 1 | 0.1 | 0.2 | 0.7 | 0.8 | ND |
| 2 | 1.6 | 2.3 | 4.3 | 0.6 | 1.1 | 17.5 | 15 | 0.2 |
| 3 | 23.7 | 14.4 | 10.4 | 3.6 | 2.9 | 54.7 | 44.9 | 1.3 |
| 4 | 34.8 | 27.3 | 20.8 | 16 | 11.8 | 22.1 | 20.9 | 10.3 |
| 5 | 25.7 | 28.4 | 25 | 36.5 | 34.3 | 5.1 | 18.9 | 59.1 |
| 6 | 12.1 | 18.3 | 21.4 | 35.5 | 39.1 | ND | 0.3 | 26.8 |
| 7 | 1.7 | 5.5 | 10.1 | 6.1 | 8 | ND | ND | 2.7 |
| 8 | 0.3 | 2.3 | 4.9 | 1.2 | 1.9 | ND | ND | 0 |
| 9 | 0.1 | 0.7 | 1.5 | 0.3 | 0.5 | ND | ND | 0 |
| 10 | 0 | 0.2 | 0.5 | 0.1 | 0.1 | ND | ND | ND |

**Table S5.** Linear regression parameters describing temporal changes in $PCB_{LN}$ (log mg/kg ww) by homolog group in each Area for quahogs (2003-2016) and scup (2003-2014). Homolog groups are denoted by number of chlorines (e.g Homolog 2 represents di-chlorobiphenyls). Red boxes denote statistical significance for the slope of $PCB_{LN}$ at alpha=0.05. Scup were sampled only in Areas 2 and 3; ns - not sampled. Trends in log-transformed $PCB_{LN}$ are displayed graphically in Figures 3c-d.

| Area | Homolog | Quahogs | | | Scup | | |
|---|---|---|---|---|---|---|---|
| | | Intercept | Slope | Slope p-value | Intercept | Slope | Slope p-value |
| 1 | 1 | -6.9394 | 0.0029 | 0.5583 | ns | ns | ns |
| | 2 | 57.3261 | -0.0286 | <0.0001 | ns | ns | ns |
| | 3 | 71.4826 | -0.0354 | <0.0001 | ns | ns | ns |
| | 4 | 60.6364 | -0.0301 | <0.0001 | ns | ns | ns |
| | 5 | 49.5989 | -0.0246 | <0.0001 | ns | ns | ns |
| | 6 | 35.7921 | -0.0179 | 0.0005 | ns | ns | ns |
| | 7 | 18.4730 | -0.0096 | 0.0280 | ns | ns | ns |
| | 8 | 8.4288 | -0.0047 | 0.2040 | ns | ns | ns |
| | 9 | 14.2987 | -0.0076 | 0.1408 | ns | ns | ns |
| | 10 | -4.5477 | 0.0017 | 0.8202 | ns | ns | ns |
| 2 | 1 | -6.2741 | 0.0026 | 0.5003 | 2.8015 | -0.0022 | 0.6509 |
| | 2 | 15.6480 | -0.0082 | 0.0165 | -3.5056 | 0.0010 | 0.9106 |
| | 3 | 57.6683 | -0.0289 | <0.0001 | -3.7878 | 0.0011 | 0.7685 |
| | 4 | 52.5674 | -0.0264 | <0.0001 | 23.9180 | -0.0125 | 0.0212 |
| | 5 | 46.0575 | -0.0231 | <0.0001 | 15.2019 | -0.0081 | 0.1593 |
| | 6 | 37.6348 | -0.0190 | <0.0001 | 5.1993 | -0.0029 | 0.6416 |
| | 7 | 8.7383 | -0.0048 | 0.0103 | 3.1276 | -0.0019 | 0.7393 |
| | 8 | -5.9781 | 0.0025 | 0.2099 | -6.0217 | 0.0024 | 0.6373 |
| | 9 | -3.1947 | 0.0011 | 0.7267 | -3.6478 | 0.0011 | 0.8113 |
| | 10 | -18.1095 | 0.0085 | 0.1736 | -2.4265 | 0.0005 | 0.9449 |
| 3 | 1 | 21.3005 | -0.0111 | 0.0171 | -3.2995 | 0.0008 | 0.8868 |
| | 2 | 0.4421 | -0.0007 | 0.8310 | -18.3097 | 0.0083 | 0.3422 |
| | 3 | 28.2092 | -0.0145 | <0.0001 | -7.0123 | 0.0027 | 0.5123 |
| | 4 | 39.0097 | -0.0198 | <0.0001 | 13.7936 | -0.0076 | 0.0626 |
| | 5 | 38.2639 | -0.0194 | <0.0001 | 22.1024 | -0.0117 | 0.0086 |
| | 6 | 25.7069 | -0.0132 | <0.0001 | 25.8190 | -0.0134 | 0.0151 |
| | 7 | 17.8268 | -0.0094 | <0.0001 | 28.8851 | -0.0149 | 0.0051 |
| | 8 | 15.0244 | -0.0080 | 0.0008 | 1.3155 | -0.0013 | 0.7657 |
| | 9 | 21.4718 | -0.0112 | 0.0031 | -4.2087 | 0.0014 | 0.7443 |
| | 10 | 21.3005 | -0.0111 | 0.0951 | -8.0544 | 0.0032 | 0.6373 |

**Table S6.** Linear regression parameters and Exposure Point Concentrations (EPCs) for $\sum PCB_{WW}$ (log mg/kg ww) in NBH quahogs and scup. Trends in log-transformed $\sum PCB_{WW}$ are shown graphically in Figure S3. EPCs for the "Present" time period represent the 95% upper confidence limit (UCL) on the mean of the three most recent years for which data are available for quahogs (2014-2016) and scup (2012-2014). All other EPCs are estimates of the geometric mean $\sum PCB_{WW}$ derived from the linear equations described herein.

| | | Quahogs | | | Scup | |
|---|---|---|---|---|---|---|
| | **Parameter/Time** | **Area 1** | **Area 2** | **Area 3** | **Area 2** | **Area 3** |
| Linear Regression Parameters | Slope | -0.0352 | -0.0343 | -0.0115 | 0.0028 | -0.0169 |
| | Intercept | 70.80 | 68.07 | 21.80 | -5.88 | 33.29 |
| | R2 | 0.2338 | 0.1546 | 0.0699 | 0.0009 | 0.0589 |
| Exposure Point Concentrations | Present | 1.4290 | 0.1380 | 0.0467 | 0.9910 | 0.4010 |
| | 1980 | 11.4604 | 1.2922 | 0.1061 | 0.3810 | 0.6760 |
| | 2015 | 0.6706 | 0.0813 | 0.0420 | 0.4759 | 0.1732 |

**Figure S1:** Concentrations in dioxin-like and non-dioxin like $\sum PCB_{LN}$ (log mg/kg ww) in quahogs (a) and scup (b) in each seafood management Area over time. Dioxin-like congeners (n=12) include all non-*ortho* (PCBs 77, 81, 126 and 169) and mono-*ortho* (PCBs 105, 114, 118, 123, 156, 157, 167 and 189) substituted PCBs capable of causing dioxin-like toxicity. All other congeners (n=124) are considered non-dioxin-like. Data are reported as annual means of $\sum PCB_{LN}$ for dioxin- and non-dioxin-like congeners measured in quahogs from each Area (n= 1 to 7 composite samples/Area/year), and in scup (n=5 composite samples/Area/year).

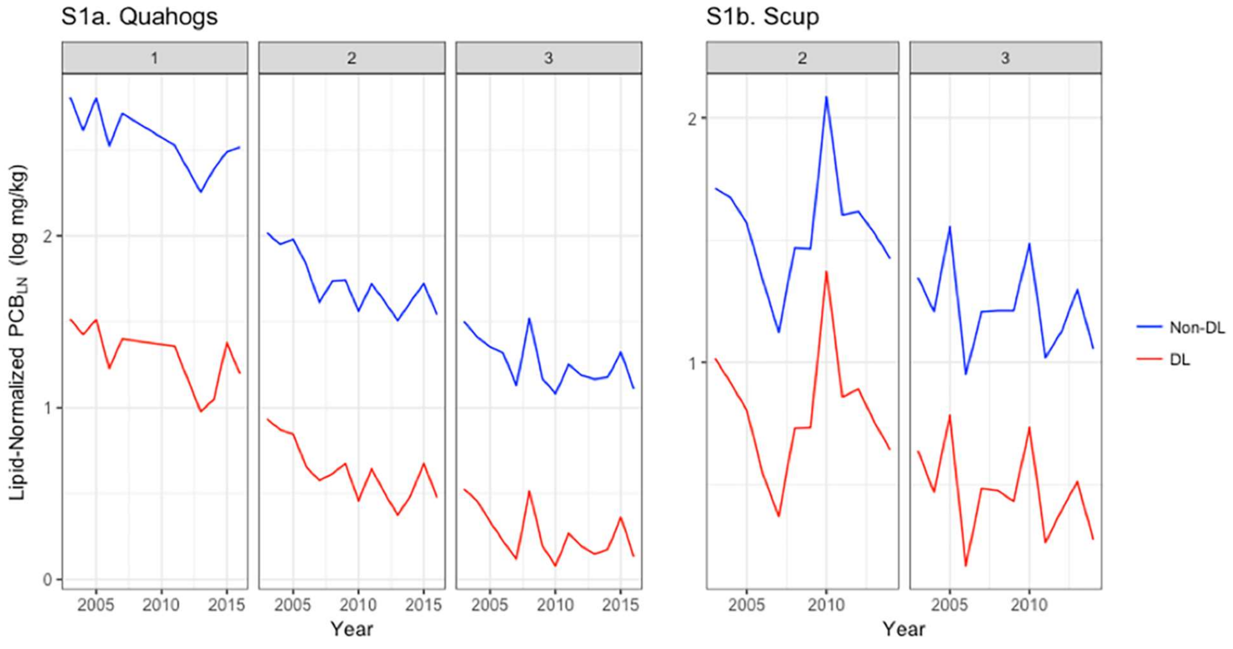

**Figure S2.** Linear regression analysis for changes in ∑$PCB_{WW}$ ((log mg/kg ww); linear regression parameters for ∑$PCB_{WW}$ (log mg/kg ww)) are presented in Table S6, and were used to estimate ∑$PCB_{WW}$ (mg/kg ww) concentrations in NBH quahogs (a) and scup (b) in 1980 and 2015. PCB estimates (mg/kg ww) at these four time points were used as Exposure Point Concentrations (EPCs) to evaluate changes in human cancer risk from PCB exposure associated with consuming NBH-harvested seafood over time. N = 1 to 7 composite samples/Area/year for quahogs and n = 5 composite samples/Area/year for scup.

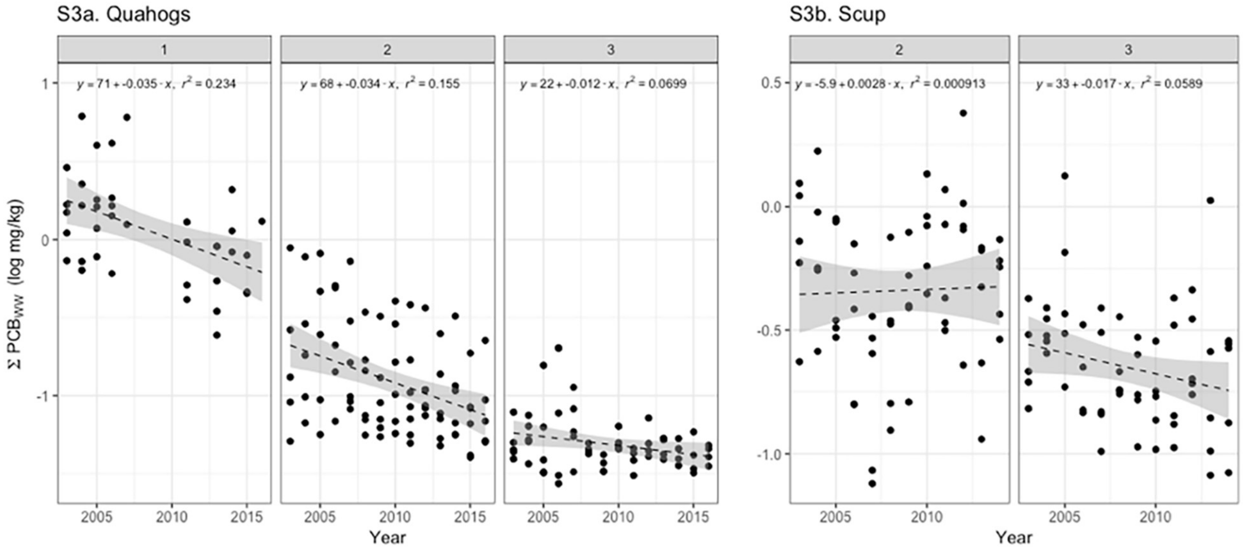
